# MCAM contributes to the establishment of cell autonomous polarity in myogenic and chondrogenic differentiation

**DOI:** 10.1101/130906

**Authors:** Artal Moreno-Fortuny, Laricia Bragg, Giulio Cossu, Urmas Roostalu

**Affiliations:** Manchester Academic Health Science Centre, Division of Extracellular Matrix and Regenerative Medicine, Faculty of Biology, Medicine and Health, University of Manchester, UK

**Keywords:** cell polarity, MCAM, myogenesis, chondrogenesis, differentiation

## Abstract

Cell polarity has a fundamental role in shaping the morphology of cells and growing tissues. Polarity is commonly thought to be established in response to extracellular signals. Here we used a minimal *in vitro* assay that enabled us to monitor the determination of cell polarity in myogenic and chondrogenic differentiation in the absence of external signalling gradients. We demonstrate that the initiation of cell polarity is regulated by melanoma cell adhesion molecule (MCAM). We found highly polarized localization of MCAM, Moesin (MSN), Scribble (SCRIB) and Van-Gogh-like 2 (VANGL2) at the distal end of elongating myotubes. Knockout of MCAM or elimination of its endocytosis motif does not impair the initiation of myogenesis or myoblast fusion, but prevents myotube elongation. MSN, SCRIB and VANGL2 remain uniformly distributed in MCAM knockout cells. We show that MCAM is also required at early stages of chondrogenic differentiation. In both myogenic and chondrogenic differentiation MCAM knockout leads to transcriptional downregulation of *Scrib* and enhanced MAP kinase activity. Our data demonstrates the importance of cell autonomous polarity in differentiation.

## INTRODUCTION

Determination of cell polarity is one of the core principles in developmental biology (Leung et al., 2016). Experiments in model organisms, ranging from the fruit fly to mouse, have demonstrated how cell polarity and differentiation are influenced by morphogen gradients and local tissue architecture (Heller and Fuchs, 2015; Lawrence and Casal, 2013; Lawrence et al., 2007). However, an important question has remained unanswered – how differentiating cells achieve polarity in the absence of external signals? An example here is the skeletal muscle cell that even *in vitro* elongates in a highly polarized orientation.

Cell polarity is established by an intricate network of positive and negative protein interactions (Campanale et al., 2017; Devenport, 2014). SCRIB (Scribble) is a tumour suppressor and one of the regulators of cell polarity. It interacts with Rho Guanine Nucleotide Exchange Factor 7 (ARHGEF7; beta-PIX), thereby controlling cytoskeletal organization and diverse signalling pathways (Audebert et al., 2004). SCRIB binds directly VANGL2 (Van-Gogh-like 2), another principal component of cell polarity establishment (Kallay et al., 2006). Additional members of the SCRIB complex include DLG1 (discs, large homolog 1) and LLGL1 (Lethal Giant Larvae Homolog 1). Knockout of these polarity regulators leads to severe embryonic malformations in mice (Caruana and Bernstein, 2001; Klezovitch et al., 2004; Murdoch et al., 2003; Murdoch et al., 2001; Pearson et al., 2011; Yin et al., 2012). Disruption of basolateral SCRIB polarity complex causes expansion of apical PAR3-PAR6-aPKC complex, illustrating reciprocally repressive interactions (Bilder and Perrimon, 2000; Bilder et al., 2003). PAR3 and 6 are PDZ domain containing scaffolding proteins that are essential in polarity establishment (Etemad-Moghadam et al., 1995; Hung and Kemphues, 1999). They interact with GTPase CDC42 (Cell Division Cycle 42) to regulate downstream signalling pathways (Joberty et al., 2000). An ever increasing number of additional proteins are involved in the establishment of cellular asymmetry. Much of our current knowledge about cell polarity arises from research in invertebrate models and only few studies have addressed its role in vertebrate mesoderm development. It has been shown how in embryonic development WNT11 can act as a directional cue for myotube elongation by activating the planar cell polarity pathway (Gros et al., 2009). Nevertheless, multiple concurrent signalling pathways may be induced by WNT11, which also regulates neuromuscular junction formation via β-catenin and VANGL2 (Messeant et al., 2017). Polarity pathway components are involved in the asymmetric division of satellite cells in the skeletal muscle (Le Grand et al., 2009; Ono et al., 2015). Cell polarity guides also chondrocyte proliferation in the elongation of long bones (Gao et al., 2011; Li and Dudley, 2009; Wang et al., 2011).

Cell migration in response to external signals relies on asymmetric distribution of surface receptors and cytoskeleton. Such transient polarization is required for providing the cell directionality and guiding the formation of protrusions. In migrating melanoma cells non-canonical WNT signalling leads to cytoskeletal rearrangement and asymmetric distribution of MCAM (Melanoma cell adhesion molecule, CD146) through its controlled endocytosis (Witze et al., 2008). MCAM is targeted to the posterior end of the migrating cell, where it forms part of the WNT5A-receptor-actin-myosin-polarity (WRAMP) structure (Witze et al., 2008). WRAMP is stably maintained during sustained periods of directional cell migration, but disbanded in cells as they pause or change direction (Connacher et al., 2017). MCAM is highly expressed in embryonic development and is maintained in postnatal skeletal muscle satellite cells and osteogenic mesenchymal stromal cells (Alexander et al., 2016; Chan et al., 2005; Li et al., 2003; Pujades et al., 2002; Sacchetti et al., 2007; Shi and Gronthos, 2003; Shih and Kurman, 1996). It is markedly upregulated in metastatic tumours (Johnson et al., 1996; Wang and Yan, 2013). Despite its high expression throughout embryonic development, still little is known on its function in cell differentiation. It can bind directly WNT1, 3 and 5 to regulate Dishevelled (DVL) and C-JUN (Jun Proto-Oncogene) phosphorylation (Ye et al., 2013). MCAM acts via NFAT (Nuclear Factor Of Activated T-Cells) and JNK (MAPK8, Mitogen-Activated Protein Kinase 8) pathways to regulate asymmetry in zebrafish and *Xenopus* embryonic development (Gao et al., 2017). Here we aimed to investigate the role of MCAM in the establishment of cell autonomous polarity in differentiating cells. We show that MCAM is asymmetrically distributed at the tip of elongating myotube, where it colocalizes with actin binding protein Moesin (MSN) and cell polarity pathway regulators VANGL2 and SCRIB. CRISPR-Cas9 mediated knockout of MCAM or deletion of its endocytosis motif leads to loss of cell polarity and failure in myotube directional elongation. MCAM knockout has a detrimental impact on early chondrogenic differentiation. Our study reveals a novel role for MCAM in regulating cell polarity in mesoderm differentiation.

## RESULTS

### Generation and initial characterization of MCAM mutant cell lines

We sought a model that would enable us to follow cellular differentiation *in vitro* from the onset of fate commitment to terminal differentiation. We chose the multipotent embryonic 10T1/2 cells as these can be induced to chondrogenic fate by recombinant BMP2 (Denker et al., 1999; Shea et al., 2003) and to myogenic fate by exposing them to testosterone after initial brief culture with 5-azacytidine (Singh et al., 2003) (Fig. 1A). We used CRISPR-Cas9 mediated genome editing, followed by FACS and single-cell cloning to generate 3 cell lines (Fig. 1B). Lines C149 and C164 were made by targeting the second exon of *Mcam* in order to achieve complete loss-of-function effect (Fig. 1B, S1A-B). Both lines carry premature STOP codons in the first N-terminal immunoglobulin domain. The MCAM protein (648 amino acid long) has a C-terminal endocytosis motif at amino acids 643-646 and since endocytosis has been proposed to regulate asymmetric MCAM trafficking in cell migration (Witze et al., 2008) we generated a third line, U125, that carries three mutations that all lead to the specific elimination of the endocytosis motif.

**Fig. 1.**
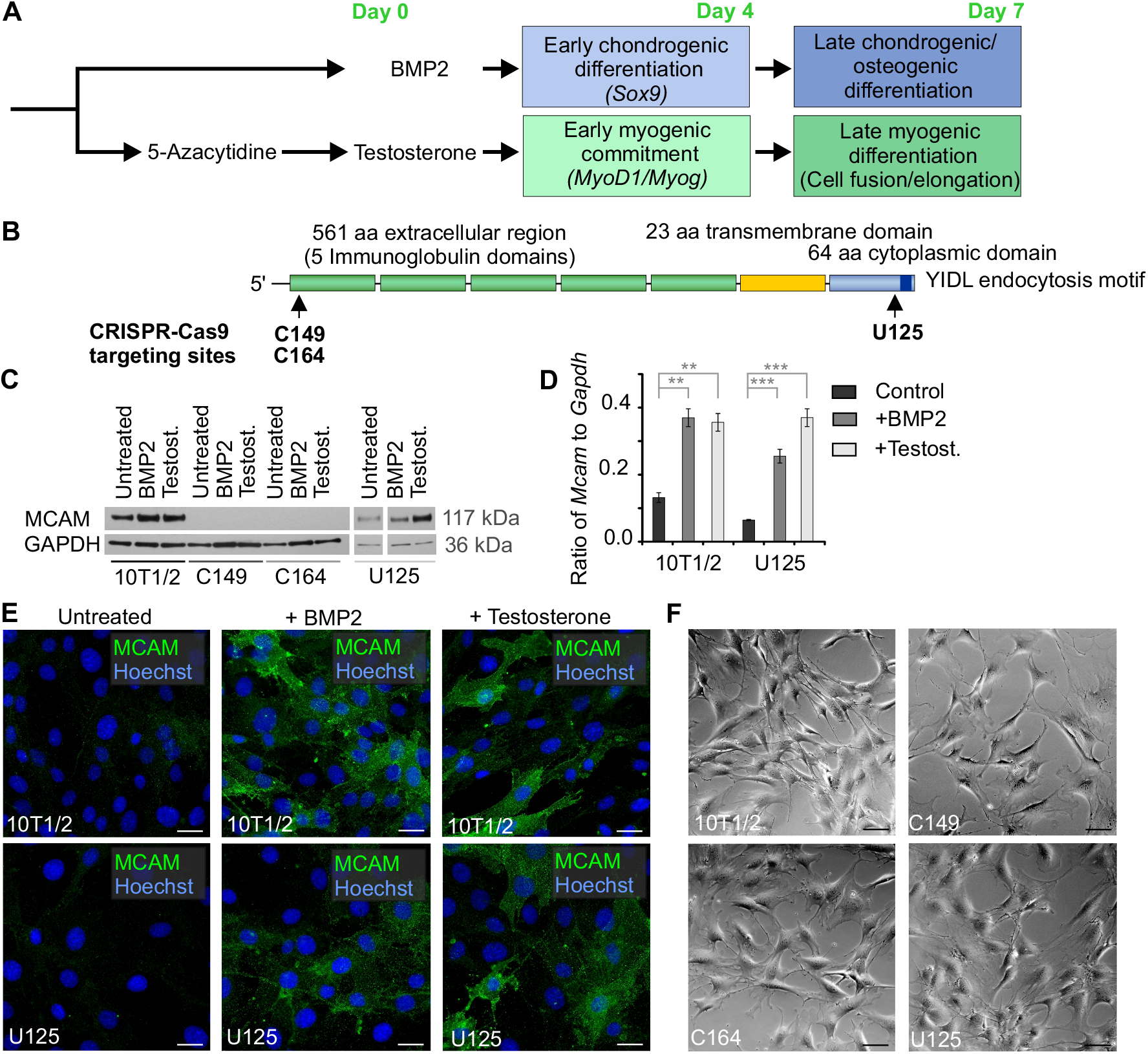
Generation of MCAM knockout cell lines. (A) Simplified diagram of the protocol used to induce myogenic and osteochondrogenic differentiation of 10T1/2 cells. (B) Schematic drawing of MCAM protein structure. CRISPR-Cas9 targeting regions for C149, C164 and U125 cell lines are indicated. Cell lines C149 and C164 both carry two mutations. C149 line has a single basepair (bp) insertion c.106_107insA and a 2 bp insertion c.106_107insCG. Both mutations lead to change of amino acid (aa) code after position 35 and STOP codon after 88 or 66 aa respectively. Line C164 has a 4 bp deletion c.103_106delCCCG and a 1 bp insertion c.106_107insT, resulting in aa code change after position 34 and 35 respectively and STOP codon after 64 and 88 aa. In line U125 3 mutations lead to elimination of the C-terminal endocytosis motif: Single bp mutation c.1899delG causes a frameshift that replaces the last 15 aa; c.1890-1908delCGGTGACAAGAGGGCTCCA affects the last 19 aa; and a larger 56 bp deletion starting from cDNA position c.1878 and encompassing 16 bp of the following intron replaces the last 22 aa of MCAM. (C) Western blot detection of MCAM in 10T1/2 cells undergoing chondrogenic (treated for 4 days with BMP2) or myogenic (treated for 4 days with testosterone) differentiation. U125 cells lack C-terminal endocytosis motif and express MCAM protein. GAPDH (Glyceraldehyde 3-phosphate dehydrogenase) – loading control. (D) RT-qPCR shows *Mcam* upregulation in 10T1/2 and U125 cells treated with either BMP2 or testosterone (mean +/- SEM; n=3; **=p<0.01; ***=p<0.001). (E) Immunocytochemistry shows MCAM upregulation in differentiating 10T1/2 and U125 cells after four day culture with BMP2 or testosterone. (F) MCAM knockout or loss of its endocytosis motif did not change the morphology of naïve fibroblasts. Scale bars: 20 μm.

*Mcam* expression was significantly upregulated in wild-type 10T1/2 and U125 cells upon differentiation, when either testosterone or BMP2 was added to the culture medium (Fig. 1C-E). Western blot analysis and immunohistochemistry proved the absence of MCAM in lines C149 and C164 (Fig. 1C, E). In their undifferentiated state the morphology of MCAM knockout cells did not differ from unedited 10T1/2 cells (Fig. 1F). Therefore MCAM is largely dispensable for fibroblasts, making our system suitable for studying its role in the context of differentiation.

### MCAM is required for chondrogenic and myogenic differentiation *in vitro*

BMP2 exposure of 10T1/2 cells leads to sequential activation of chondrogenic and osteogenic differentiation programs (Shea et al., 2003). BMP2 induced chondrogenic differentiation of 10T1/2 cells by 4 days in culture, which was evident in intense alcian blue staining (Fig. 2A, S2A) and upregulation of *Sox9* (Fig. 2C) (Bi et al., 1999). Knockout of MCAM abolished chondrogenic differentiation in all the established cell lines (Fig. 2A, C). Wild-type BMP2 treated cells upregulated hypertrophic chondrocyte marker *Col10a1* by the 7^th^ day in culture, whereas its levels remained low in MCAM mutant cells (Fig. 2D). Alkaline phosphatase positive osteogenic nodules formed in the culture of wild-type cells (Fig. 2B) and terminal osteogenic differentiation marker osteomodulin (*Omd*) was upregulated (Fig. 2E). Osteogenic differentiation was significantly impaired in the absence of MCAM (Fig. 2B, E).

**Fig. 2.**
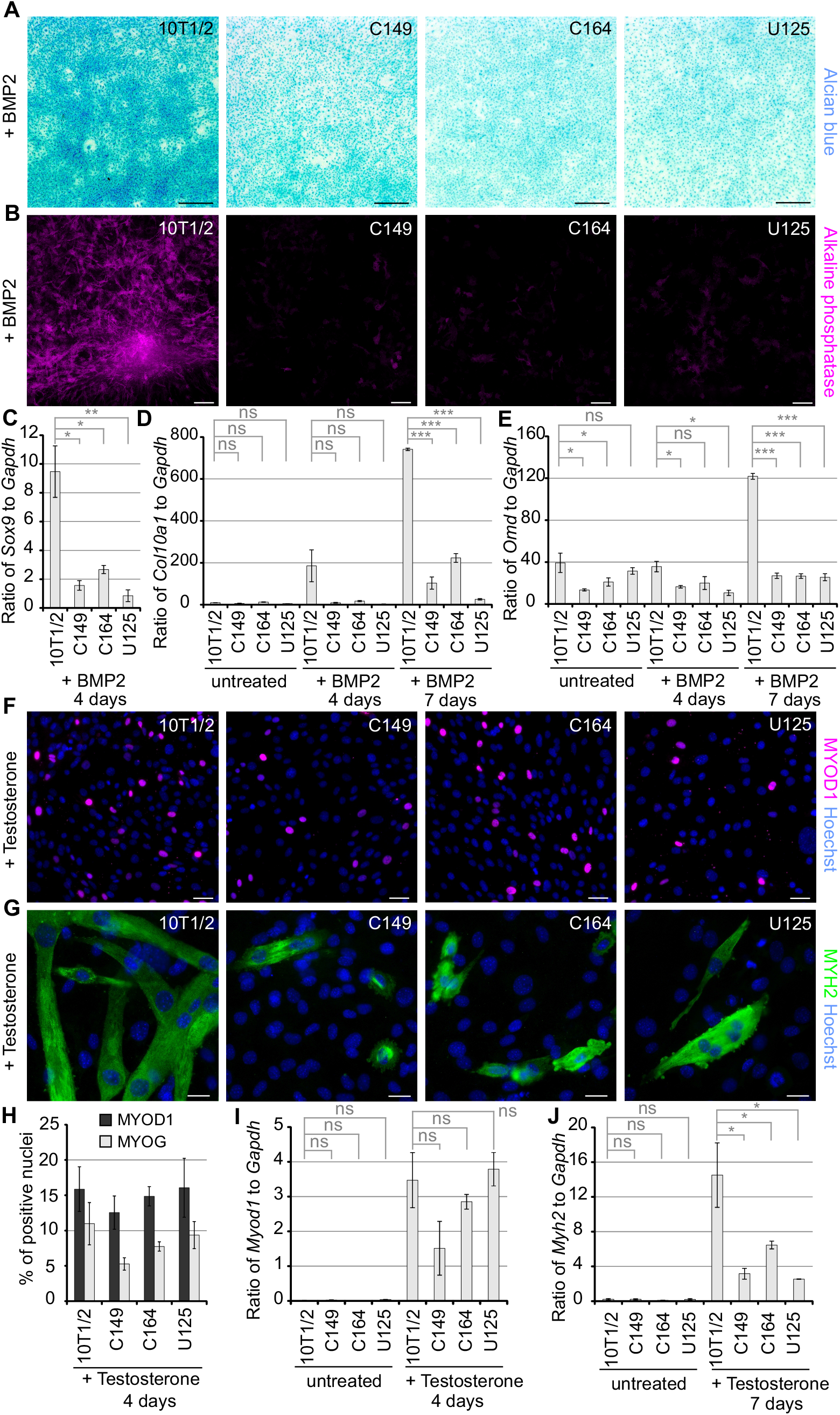
Loss of MCAM function impairs chondrogenic and myogenic differentiation. (A) Alcian blue staining showing that in comparison to wild-type 10T1/2 cells chondrogenic differentiation fails in MCAM mutant cells (C146, C164, U125) treated for 4 days with BMP2 (blue staining corresponds to chondrogenic differentiation). (B) MCAM loss of function abolished the formation of alkaline phosphatase positive osteogenic nodules in cultures treated for 7 days with BMP2. RT-qPCR (mean +/- SEM; n=3; ns - not significant; *=p<0.05; **=p<0.01; ***=p<0.001) analysis revealed (C) no upregulation of *Sox9* and significantly diminished expression of (D) *Col10a1* and (E) *Omd* in MCAM knockout cell lines treated with BMP2. (F) MCAM knockout did not prevent MYOD1 upregulation after 4 day treatment with testosterone. (G) 7 day exposure to testosterone triggered extensive formation of multinucleated myotubes in wild-type 10T1/2 cells, whereas MCAM knockout cells failed to elongate into myotubes. (H) Quantification of MYOD1 and myogenin (MYOG1) positive nuclei after 4 day exposure to testosterone (no significant differences detected; t-test; n=5 independent cell culture experiments). RT-qPCR analysis of (I) *MyoD1* expression showing testosterone induced upregulation in both wild-type and MCAM knockout cells at 4 days, and (J) significantly reduced expression of myosin heavy chain (*Myh2*) in *Mcam* mutant cells at 7 days. Scale bars: A, 500 μm; B, 200 μm; F, 50 μm; G, 20 μm.

In myogenic promoting conditions early transcriptional regulators *Myod1* and myogenin (*Myog*) were induced in both 10T1/2 and *Mcam* mutant cells after 4 days exposure to testosterone (Fig. 2F, H-I, S2B-C). Despite this seemingly unaffected initiation of the myogenic program, terminal differentiation into multinucleated myotubes was severely impaired in cells lacking MCAM or its endocytosis motif. MCAM mutant cells fused into multinucleated myogenic cells that failed to elongate (Fig. 2G, S2D). There was a correspondingly reduced expression of myosin heavy chain in *Mcam* mutant cell lines (Fig. 2J). These data show the importance of MCAM in regulating mesodermal differentiation. Our results highlight the differences between the role of MCAM in chondrogenic and myogenic commitment, whereby it is essential at the onset of the former, but not until the elongation stage in the latter.

We next analysed whether loss of MCAM may lead to enhanced expression of other CAMs. We found that neither neural cell adhesion molecule (*Ncam1*) isoforms 1 and 3 were upregulated in the knockout cell lines in comparison to wild-type cells (Fig. 3A). As expected, *Ncam1* skeletal muscle specific isoform was significantly upregulated in wild-type 10T1/2 cells in comparison to knockout cells after 7 day exposure to testosterone. Intercellular adhesion molecule 1 (*Icam1*) was upregulated significantly only in wild-type cells in myogenic growth conditions (Fig. 3B). *Icam2* was expressed at low levels in all the cell lines. Vascular cell adhesion molecule 1 (*Vcam1*) was most highly expressed in undifferentiated cells, but its expression levels did not depend on the presence of MCAM (Fig. 3C). These data indicate that loss of MCAM does not cause a compensatory upregulation of other CAMs and rather the absence of MCAM leads to downregulation of CAMs that are characteristic of differentiating cells.

**Fig. 3.**
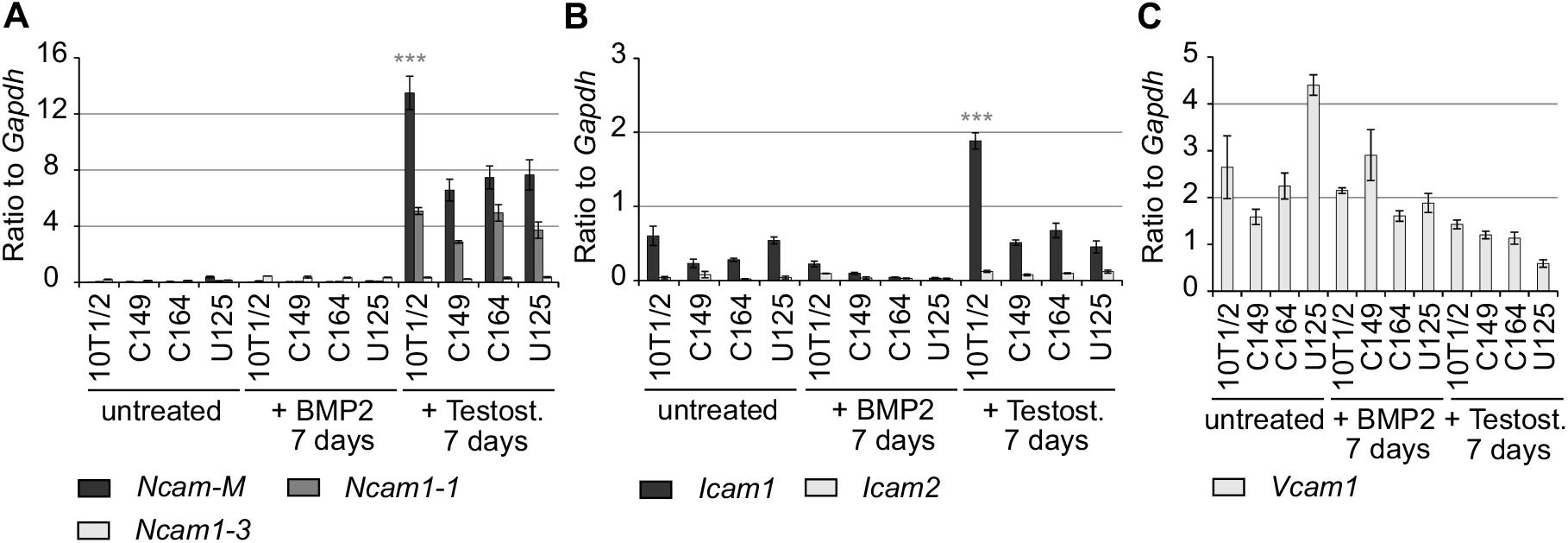
MCAM knockout effect on cell adhesion molecule expression. (A) Expression of *Ncam1* isoforms 1 and 3 and muscle specific isoform *Ncam1-M* relative to *Gapdh*. Untreated cells and cells treated for 7 days with BMP2 or testosterone. *Ncam1-M* was significantly (P<0.001; n=3) induced in wild-type cells exposed for 7 days to myogenic growth conditions in comparison to MCAM knockout cell lines. (B) Expression levels of *Icam1* and 2. *Icam1* was significantly upregulated (p<0.001; n=3) in wild-type cells after 7 day exposure to testosterone in comparison to MCAM knockout cell lines. (C) Expression levels of *Vcam1* did not significantly differ among the studied cell lines.

### Asymmetric distribution of MCAM regulates cell autonomous polarity

We next addressed the mechanisms by which MCAM influences differentiation. First we approached the established signalling pathways downstream of MCAM: it is known to induce DVL2 and C-JUN phosphorylation in cancer cells and its knockdown leads to the induction of canonical WNT pathway (Ye et al., 2013). We noticed increased C-JUN phosphorylation under both osteochondrogenic and myogenic conditions, but this occurred even in the absence of MCAM (Fig. 4A). DVL2 activity decreases in myogenesis (Yamaguchi et al., 2012) and we confirmed this here, however there were no consistent differences between wild-type and MCAM mutant cells (Fig. 4B). We did not detect significant changes in the expression of canonical WNT transcription factors T-cell factor 4 (*Tcf4*) and Lymphoid Enhancer Binding Factor 1 (*Lef1)* (Fig. S3A). Neither did we observe nuclear translocation of β-catenin or change in WNT5A staining (Fig. S3B-C). In cancer cells MCAM has been linked with *Id1* expression (Zigler et al., 2011), which is a known inhibitor of myogenesis (Jen et al., 1992). However, we did not detect altered *Id1* expression in MCAM mutant cell lines, indicating that this pathway is unlikely to explain defective myogenic differentiation (Fig. 4C).

**Fig. 4.**
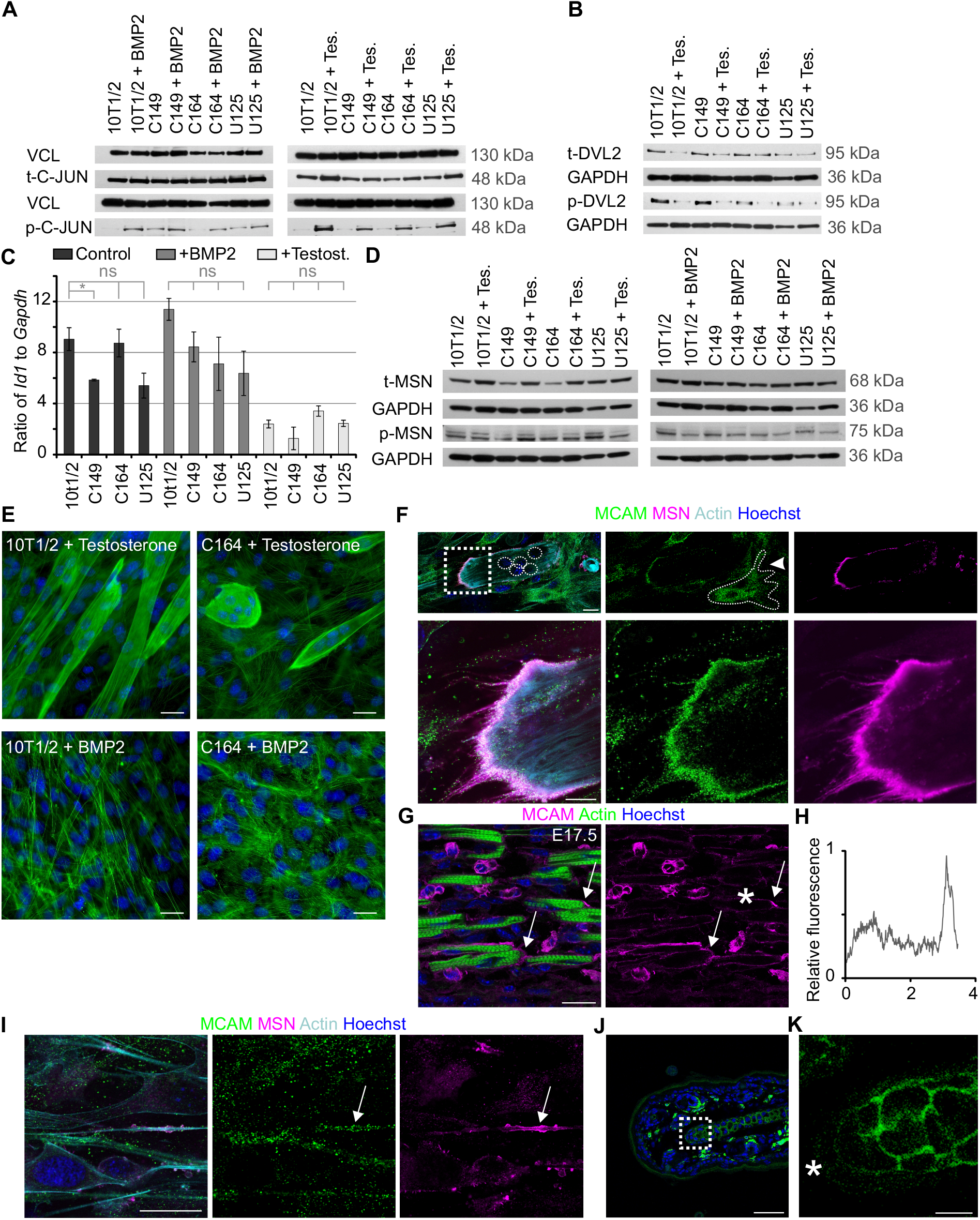
Asymmetric distribution of MCAM and MSN in differentiating cells. (A) MCAM loss of function did not lead to changes in total C-JUN level (t-C-JUN). Differentiation induced phosphorylation of C-JUN (p-C-JUN), which occurred independently of MCAM. VCL (Vinculin) - loading control. (B) Myogenic differentiation led to downregulation of total DVL2 and its phosphorylated form (p-DVL2), yet this occurred regardless of MCAM activity. GAPDH – loading control. (C) RT-qPCR showing downregulation of *Id1* expression (mean +/- SEM; n=3; ns - not significant; *=p<0.05) in cells undergoing myogenic differentiation. (D) Western blot analysis of total (t-) and phosphorylated (p-) MSN after 7 day exposure to BMP2 or testosterone. (E) Actin staining (Phalloidin-Alexa-488) illustrates large morphological changes in MCAM mutant cells. Multinucleated myotubes fail to elongate in the absence of MCAM (upper panel, 7 day treatment with testosterone) and chondrogenic cells show impaired cytoskeletal assembly (7 day exposure to BMP2). (F) In multinucleated myotubes MCAM colocalizes with MSN in the distal end of the cell, often in cytoplasmic processes. In undifferentiated cells MCAM is primarily localized to cytoplasmic vesicles (arrowhead, outlined). Upper panel illustrates low magnification image of a multinucleated myotube (nuclei highlighted in circles; single confocal plane). Magnified image of the boxed area is shown underneath. (G) In foetal mouse limbs (E17.5) MCAM can be seen at ends of differentiating myotubes. Single confocal plane is shown. (H) Quantification of MCAM staining intensity over a single myotube indicated with asterisk on figure G. MCAM shows enrichment at one end of the myotube. (I) MCAM colocalizes with MSN also in 10T1/2 cells exposed for 7 days to BMP2. (J-K) MCAM localization in juvenile (P20) mouse ear cartilage. (K) Magnification of the boxed area. At the edge of the growing cartilage (asterisk) MCAM staining is weak and cytoplasmic, but acquires membrane localization in mature cartilage. Single confocal plane is shown. Scale bars: E-G, I, 25 μm; J, 50 μm; K, 10 μm.

Several lines of evidence link MCAM with actin cytoskeleton. It interacts directly with actin binding proteins hShroom1 (Dye et al., 2009) and MSN (Luo et al., 2012). We tested the phosphorylation levels of MSN, but detected only a small decrease in its activity in MCAM mutant cells (Fig. 4D). We found significant morphological changes in MCAM mutant cells, which are clearly visible in actin cytoskeletal organization (Fig. 4E). Most notably, myoblasts fused together into multinucleated cells that however expanded in size without elongating and aligning the actin cytoskeleton along the direction of the cell. This led us to hypothesize that myotube elongation relies on cell autonomous initiation of polarity and defects in this process may underlie the phenotype seen in MCAM mutant cells. Furthermore, MSN is a known regulator cytoskeletal organization in cell polarity (Fehon et al., 2010; Jankovics et al., 2002; Medina et al., 2002; Polesello et al., 2002; Speck et al., 2003). Abnormal localization of polarity complex rather than large changes in its activity may therefore lead to the observed differentiation defects. We analysed whether MCAM and MSN are asymmetrically distributed in differentiating cells. We found that in elongating myotubes MCAM colocalizes with MSN at the distal tip of the cell (Fig. 4F). We detected asymmetric distribution of MCAM also *in vivo*, in E17.5 mouse forelimb developing myotubes (Fig. 4G, H, S3D). Polarity in chondrogenic differentiation remains less obvious, although we could detect MCAM and MSN colocalization in 10T1/2 cells exposed to BMP2 (Fig. 4I). We analysed growing juvenile mouse ear cartilage and found that at the edges, where the cartilage is growing, MCAM is weakly expressed and cytoplasmic, but acquires membrane localization in mature cartilage (Fig. 4J-K). Polarization may occur transiently in cartilage maturation and remains difficult to establish *in vivo*.

We next analysed the localization of cell polarity associated proteins in MCAM mutant cells in comparison to wild-type cells undergoing myogenic differentiation. Similarly to MSN and MCAM we found highly asymmetric distribution of VANGL2 at the distal tips of elongating myotubes (Fig. 5A-B, S4A). Remarkably, in MCAM knockout cells such polarity was not established and MSN labelled the whole plasma membrane, whereas VANGL2 was uniformly distributed across the cell (Fig. 5C, S4A). We detected colocalization of SCRIB with MCAM at the tip of an elongating myotube (Fig. 5D, S4B). Knockout of MCAM led to uniform distribution of SCRIB (Fig. 5E) and importantly, also its transcriptional downregulation (Fig. 5H). SCRIB gene regulatory elements are not known and the precise transcriptional link remains to be established. Intriguingly, elimination of MCAM endocytosis motif led to frequent localization of SCRIB on actin filaments (Fig. S4C-C’’). SCRIB trafficking to cortical actin was recently established to rely on β spectrin (Boeda and Etienne-Manneville, 2015). Our data suggests that MCAM participates in SCRIB trafficking.

Since SCRIB complex antagonizes cell cortex targeting of PAR3, we analysed its localization and found that in wild-type cells PAR3 was largely cytoplasmic, whereas in MCAM mutant myotubes it was frequently enriched at the membrane (Fig. 5F-G). Also in MCAM endocytosis motif mutant cells, VANGL2 and PAR3 were mislocalized (Fig. 5I). We conclude that subcellular trafficking of MCAM is important in the establishment of cell autonomous polarity in myogenic differentiation.

**Fig. 5.**
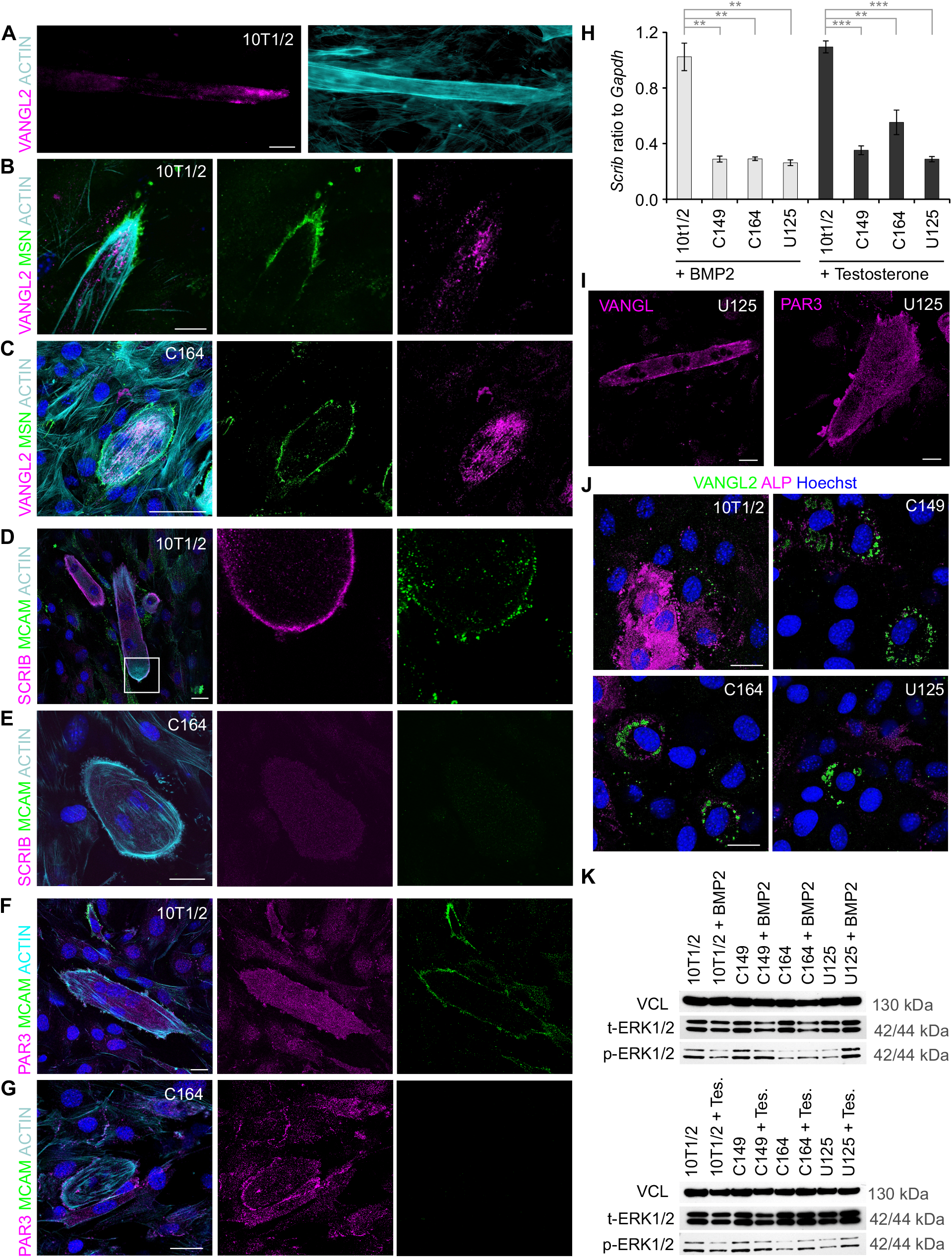
MCAM is required to establish cell autonomous polarity. (A) In elongating myotubes (10T1/2 cells treated with testosterone for 7 days) VANGL2 is localized asymmetrically at the tip of the cell. (B) The VANGL2 enriched tip of the cell is marked by MSN. (C) In MCAM knockout C164 cells myotube elongation fails, MSN labels the whole plasma membrane and VANGL2 is spread across the cytoplasm. (D) Highly polarized localization of MCAM and SCRIB at the distal end of growing wild-type myotube. (E) In MCAM knockout cells SCRIB levels remain low and it is spread evenly in the cell. (F) In wild-type myotubes PAR3 remains cytoplasmic, whereas (G) in MCAM knockout C164 cells it can be detected at the cell cortex. (H) RT-qPCR demonstrates reduced expression of *Scrib* in MCAM mutant cell lines. Cells were treated for 7 days with BMP2 or testosterone (n=3; **=p<0.01; ***=p<0.001). (I) Deletion of MCAM endocytosis motif leads to similar polarity defects as complete MCAM elimination. VANGL2 is evenly spread in U125 cells and PAR3 accumulates in cell cortex. (J) In chondrogenic differentiation VANGL2 was observed asymmetrically in limited number of cells. In MCAM mutant cell lines (C149, C164, U125) VANGL2 accumulated around the nucleus. (K) Initiation of myogenic (4 day culture with testosterone) and chondrogenic differentiation (4 day culture with BMP2) led to downregulation of ERK1/2 phosphorylation (p-ERK1/2). Instead in MCAM mutant cell lines ERK1/2 phosphorylation increased. Scale bars: 25 μm.

We found significant downregulation of *Scrib* in MCAM mutant cell lines in osteochondrogenic differentiation (Fig. 5H). Intriguingly, while in wild-type BMP2 exposed cells VANGL2 showed only limited asymmetric localization, in the absence of MCAM it accumulated across the cytoplasm (Fig. 5J). We could also detect membrane targeting of PAR3 in MCAM mutant cells, although it remained less prominent than in myogenic differentiation (Fig. S4D). We propose that cell polarity has a more dynamic role in chondrogenic differentiation.

Several signalling pathways are regulated by cell polarity. Loss of SCRIB leads to enhanced MAP kinase pathway (ERK1/2) activity (Nagasaka et al., 2010) and since tight control over ERK1/2 signalling is required in chondrogenesis (Bobick and Kulyk, 2004) we tested its activity in MCAM mutant cell lines. While in both chondrogenic and myogenic differentiation wild-type cells downregulate ERK1/2 phosphorylation, in MCAM mutant cells their levels remained as high as in undifferentiated cells, providing a mechanistic insight into signalling changes in the cells (Fig. 5K).

## DISCUSSION

We have established an *in vitro* assay that enabled us to monitor cell polarity determination throughout myogenic and chondrogenic differentiation. Our data enables us to propose a model on how myotubes achieve elongated tubular morphology. We propose that highly asymmetric localization of MCAM enables the adhesion and anchoring of the rear end of an elongating myotube (Fig. 6). MCAM is known to interact directly with MSN (Luo et al., 2012) and this interaction explains how MCAM dependent polarity is translated to actin cytoskeleton arrangement in an elongating cell. We found that MCAM is required for the asymmetric distribution of cell polarity regulators SCRIB and VANGL2. In the absence of MCAM asymmetric localization of MSN fails and actin cytoskeleton is dispersed in diverse directions across the cell, preventing myotube elongation. Thus the emerging model suggests that myotube elongation is reminiscent of cancer cell or leukocyte migration, with the exception that the rear of the cell is firmly adhered and does not follow the leading edge. The cell polarity associated proteins have well established roles in cell migration (Ebnet et al., 2017). Both MSN and MCAM are highly expressed in metastatic melanoma cells and known to regulate cell migration (Connacher et al., 2017; Estecha et al., 2009; Lei et al., 2015; Witze et al., 2008). Current evidence indicates that MCAM is a non-selective receptor for a wide number of extracellular signalling molecules that include WNT5A, WNT1, FGF4 and S100A8/A9 (Gao et al., 2017; Ruma et al., 2016; Ye et al., 2013). It is also a co-receptor for vascular endothelial growth factor receptor 2 (Jiang et al., 2012). Therefore diverse signals are capable of inducing or modulating MCAM activity *in vivo*. In our *in vitro* system cells polarized in the absence of extracellular signalling molecule gradients, suggesting that cells have also an autonomous polarity-based mechanism that regulates their shape. This intrinsic mechanism may explain how cell polarity and distinct morphology can arise in tissues such as skeletal muscle and cartilage where, due to high density and consequent poor penetrance, morphogen gradients are difficult to be established. Similarly to our study, merlin and ERM-family proteins (ezrin, radixin, MSN) were found capable of inducing cell cortex cytoskeleton asymmetry in the absence of external cues in epithelial cells (Hebert et al., 2012).

**Fig. 6.**
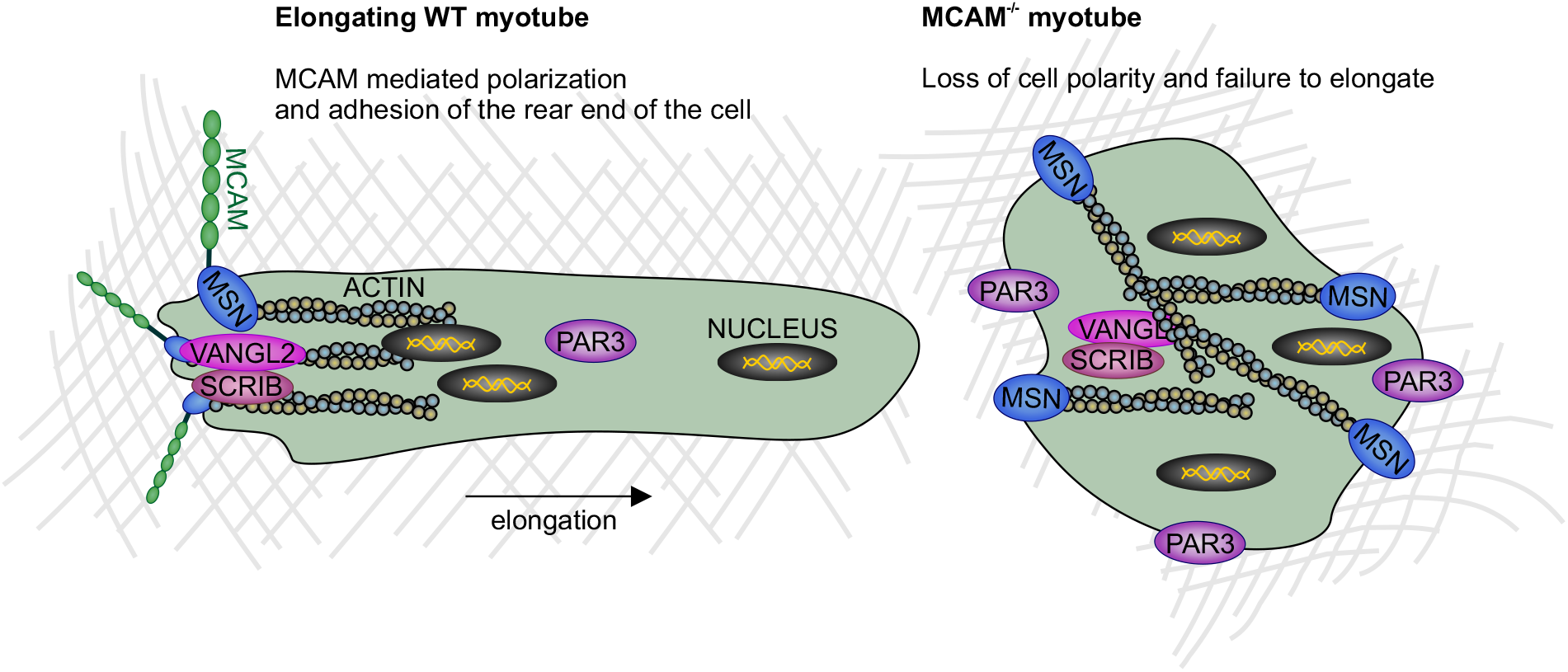
Model for molecular mechanisms regulating myotube elongation. MCAM is transiently enriched at one end of an elongating myotube where it enables cell adhesion to the extracellular matrix. The opposite leading edge can move forward to lengthen the cell. MCAM interaction with MSN enables directed orientation of actin cytoskeleton in the direction of cell elongation plane. Polarity pathway components SCRIB and VANGL2 colocalize with MCAM, whereas PAR3 is retained in the cytoplasm. In the absence of MCAM MSN loses its asymmetric distribution and consequently actin cytoskeleton fails to be polarized. In MCAM knockout cells SCRIB and VANGL2 lose asymmetric distribution and PAR3 is targeted to broad areas in the cell cortex.

The importance of cell polarity has remained poorly characterized in cartilage. Non-canonical WNT signalling controls cell division plane in the development of long bones (Li and Dudley, 2009). Loss of VANGL2 leads to limb defects in mice (Wang et al., 2011). We show that MCAM knockout leads to failure to initiate chondrogenic differentiation program *in vitro* and causes mislocalization of actin cytoskeleton and VANGL2. We hypothesize that in the absence of MCAM tight intercellular contacts that are needed in chondrogenesis even *in vitro*, fail to be established. It is likely that the cell autonomous polarity based mechanism is very transient. Future live imaging studies are needed to determine the precise role of individual proteins in this process and their dynamic interactions. Mice deficient of MCAM or MSN do not show significant developmental defects, albeit in both models abnormalities have been observed in postnatal pathological conditions (Doi et al., 1999; Jouve et al., 2015; Okayama et al., 2008; Tu et al., 2013; Zeng et al., 2014). ERM proteins show functional redundancy in most cell types, explaining the limited phenotype of MSN knockout mice (Fehon et al., 2010). It is likely that *in vivo* other proteins may compensate for the lack of MCAM. The hereby used *in vitro* system has enabled us to establish its role in a minimal environment with limited ligands and adhesion substrates. Our data collectively provides evidence that cellular differentiation relies on an intrinsic, adhesion receptor dependent polarity mechanism.

## ACKNOWLEDGEMENTS

U.R. was supported by BBSRC Anniversary Future Leader Fellowship (BB/M013170/1). G.C. was supported by BHF (PG/14/1/30549), MRC (MR/P016006/1), Duchenne Parent Project (Italy) and Fundació La Marató grants. We are grateful to M. C. Jackson (Flow Cytometry Core Facility, Faculty of Biology, Medicine and Health) for help with cell sorting and to A. Adamson for advice on genome editing.

## AUTHOR CONTRIBUTIONS

All authors discussed the results, analysed data and wrote the manuscript. A.M.F. conducted gene expression experiments, L.B. carried out protein analysis, G.C. discussed results and contributed to the manuscript and U.R. designed the study, generated cell lines and carried out microscopy experiments.

## MATERIALS AND METHODS

### Genome editing

CRISPR-Cas9 guide RNAs were designed in Benchling software (Benchling, Inc.). For N-terminal mutations in cell lines C149 and C164 guide RNA GCCAGTACCCACGCCCGACC/**TGG** and for the C-terminal mutation in line U125 guide RNA GGGCAGCAACGGTGACAAGA/**GGG** was used (PAM sequence in bold). These guides were cloned into SpCas9(BB)-2A-GFP plasmid as previously described (Ran et al., 2013). pSpCas9(BB)-2A-GFP was a gift from Feng Zhang (Addgene plasmid # 48138). Authenticated 10T1/2 cells were purchased from Public Health England and cultured in MEM (Gibco), supplemented with 1% L-Glut, 10% FBS, 1% PS, 1x MEM Non-Essential Amino Acids Solution. Cells were transfected with guide RNA containing plasmid using Lipofectamine 3000 (Thermo Scientific) and after 24 h culture the GFP expressing cells were sorted into 96 well plates by BD FACS Aria. Confluent wells were split to isolate genomic DNA. The guide RNAs were designed so that indels would destroy a restriction enzyme binding site (XcmI for N-terminal and BpmI for Ctermina edit). The PCR product was further verified by Sanger sequencing and analysis using CRISPR-ID (Dehairs et al., 2016). Selected lines were confirmed by cloning PCR product and sequencing individual bacterial colonies. For differentiation assays cells were plated at a density of 3200/cm^2^. Myogenic differentiation of 10T1/2 cells was carried out as described (Singh et al., 2003), by initial 3-day culture in 20 μM 5-Azacytidine (Sigma Aldrich, A2385), followed by 3 day recovery in growth medium and 4-7 day incubation with 100 nM testosterone. For chondrogenic differentiation cells were incubated with recombinant BMP2 (200 ng/ml, R&D, 355-BM-050) for 4-7 days.

### Alcian blue and alkaline phosphatase staining

Alcian Blue staining was carried out by washing the sample 3×5 minutes in PBS supplemented with 0.1% Tween (PTW) and 2×5 minutes in 3% Acetic acid. Thereafter Alcian Blue (Sigma-Aldrich, 1% solution in 3% Acetic acid) was added to the sample for 30 minutes, after which it was washed off with 3% Acetic acid followed by 3 washed in PBS (5 minutes each). Alkaline phosphatase staining was carried out using Vector Laboratories Red AP kit.

### Immunohistochemistry

Cells were grown on glass-bottom plates (24-well Sensoplate, Greiner Bio-One), washed with PBS and fixed in 4% paraformaldehyde for 10 minutes. The fixed cells were washed 3×5 minutes in PTW and incubated for 3 h in blocking buffer (10% donkey serum in PTW) and overnight with primary antibodies in blocking buffer. Second day the samples were washed 3×5 minutes and 3×30 minutes in PTW, incubated for 30 minutes in 1% BSA in PTW and for 2 h with secondary antibodies in PTW (room temperature). Following the removal of the secondary antibody the samples were washed 3×15 minutes in PTW and 3×15 minutes in PBS. Inverted Zeiss Axio Observer Z1 was used together with Zeiss Zen software. Leica SP5 confocal was used together with the Leica Application Suite software. Images were acquired at 2 times line average and 3-5 times frame average, using sequential scanning mode. Single focal planes are shown.

### Western blot

Cell samples were aspirated of media, washed 3 times with cold PBS then placed onto ice. Cold lysis (RIPA) buffer was added and the cells were scraped into sterile, pre-cooled tubes. These were incubated (with rotation) for 30-60 min at 4°C. After incubation, the samples were centrifuged and supernatant removed to a new pre-cooled tube. Protein concentration of each sample was determined using a Bradford assay. 20 ng of each sample was reduced and ran on a 4-12% NuPage Novex precast gel, following the NuPage Denaturing Electrophoresis protocol. The gels were transferred onto nitrocellulose membranes following standard protocol. The membranes were blocked using either 5% non-fat milk or 5% BSA (for phosphorylation-specific antibodies) in TBS-Tween for 60-120 min at RT. Primary antibodies were added into the relevant blocking solution and membranes incubated overnight at 4°C. After incubation they were washed 3× with TBS-Tween. Secondary antibodies (in fresh blocking solution) were added to the membranes and incubated for 60-120 min at RT. They were washed 3× with TBS-Tween after incubation. HRP substrate was added to each membrane. Chemiluminescent film was exposed to the membrane and developed using an automatic processor, JP-33.

### Antibodies

The antibodies were used at the following concentrations: Rat anti-MCAM (R&D MAB7718; ICC 1:100), Sheep anti-MCAM (R&D AF6106; WB 1:1000), Goat anti-WNT5A (R&D AF645; ICC 1:50), Rabbit anti-LEF1 (Cell Signalling 2230; 1:100), Mouse anti-MyoD1 (Dako Clone 5.8A; ICC 1:200), Mouse anti-MYOG (The Developmental Studies Hybridoma Bank F5D; ICC 1:2), Mouse anti-MYH (The Developmental Studies Hybridoma Bank MF20; ICC 1:2), Rabbit anti-C-JUN (Cell Signalling 9165; WB 1:500), Rabbit anti-Phospho-c-JUN (Ser73) (Cell Signalling 3270; WB 1:500), Goat anti-VCL (Santa Cruz sc-7649; WB 1:2000), Mouse anti-GAPDH (Abcam ab125247; WB 1:2000), Rabbit anti-DVL2 (Abcam ab22616; WB 1:1000), Rabbit anti-phospho-DVL2 (Abcam ab124933; WB 1:1000), Rabbit anti-MSN (Abcam ab52490; ICC 1:100, WB 1:5000), Rabbit antiphospho-MSN (Abcam ab177943; WB 1:500), Goat anti-VANGL2 (Santa Cruz sc-46561; ICC 1:100), Rabbit anti-SCRIB (Biorbyt orb337106; ICC 1:200), Rabbit anti-PAR3 (Novus Biologicals NBP1-88861; ICC 1:200), Rabbit anti ERK1/2 (Cell Signaling 4695; WB 1:1000), Rabbit antiphospho-ERK1/2 (Cell Signaling 4370; WB 1:2000), Phalloidin-488 (Thermo Fischer A12379, ICC 1:100). Diverse secondary Alexa dye conjugated antibodies were purchased from Life Technologies/Thermo Scientific and used at 1:500 dilution. HRP conjugated secondary antibodies (Dako Donkey anti-goat P0449; Dako Goat anti-rabbit P0448; Dako Rabbit anti-mouse P0260; Life Technologies Donkey anti-sheep A16047) for Western blot were used at 1:2000 dilution. Abbreviations: ICC – Immunocytochemistry; WB – Western blot.

### RNA isolation and RT-qPCR

RNA extraction was performed using TRIzol reagent following the standard manufacturer’s procedure (Thermo Fisher Scientific). Once resuspended, RNA was quantified with Nanodrop 2000. After treatment with DNase a standardized amount of RNA was used for retro-transcritpiton using random hexamer primers (RevertAid First Strand cDNA Synthesis Kit Thermo Fisher Scientific, 1621). RT-qPCR was carried out using FastStart Essential DNA Master Mix (Roche) on a Roche LightCycler 96 system. RT-qPCRs were performed in triplicate for each sample and 3 biological replicates were used. Absolute quantification of each target was performed using a standard curve as a reference in Roche LightCycler software version 1.5. Primer sequences were acquired from Harvard PrimerBank (Spandidos et al., 2008; Spandidos et al., 2010; Wang and Seed, 2003): *Col10a1* (6753480a1), *Gapdh* (6679937a1), *Icam1* (21389311a1), *Icam2* (6754274a1), Lef1 (27735019a1), *Mcam* (10566955a1), *MyoD1* (6996932a1), *Myh2* (21489941a1), *Ncam1.1* (817984a1), *Ncam1-M* (3980168a1), *Ncam1.3* (817988a1), *Omd* (6754934a1), *Scrib* (20373163a1), Sox9 (31543761a1), *Tcf4* (7305551a1), *Vcam1* (31981430a1).

### Statistical analysis

Statistical significance of RT-qPCR results was determined by two-tailed t-test in Microsoft Excel.

